# Dissecting the Role of Flagellar Subunits in *C. difficile* Mucosal Colonization

**DOI:** 10.1101/2025.01.02.631138

**Authors:** Baishakhi Biwas, Thi Van Thanh Do, Jennifer Auchtung, Kurt H. Piepenbrink

## Abstract

*Clostridioides difficile* is a common cause of acute gastrointestinal (GI) inflammation in mammals, which can have detrimental effects on host health. *C. difficile* associated disease (CDAD) requires the secretion of high-molecular weight toxins after colonization of the GI tract. The molecular mechanisms of GI colonization by *C. difficile*, include potential interactions with host cells and the mucus layer formed from secreted mucin glycoproteins. *C. difficile* associates with the mucus layer *in vivo* and will associate with both epithelial cells and mucosal surfaces *in vitro*. Previously, we found a substantial defect in binding to mucosal surfaces for mutants of the major flagellar subunit, *fliC*, while mutation of the major subunit of type IV pili, *pilA1*, showed increased adhesion. To elucidate the mechanisms by which *C. difficile* interacts with *ex vivo* mucosal surfaces, we have measured swimming motility, mucosal adhesion and levels of flagellation by transmission electron microscopy for mutants of flagellar and T4P genes in *C. difficile* R20291. We discovered that the *pilA1* mutant showed increased flagellation, while decreases in flagellation were found for *fliC, fliD,* and *flg-*OFF (a phase-locked mutant with low transcription of the F3 flagellar operon) which were associated with both low swimming motility and low adhesion to mucosal surfaces. However, the reversed *flg-*ON mutant showed increased flagellation without a significant increase in adhesion. We also found that the *fliC* mutant was defective in binding to mucus-secreting HT-29 MTX cells, but not HT-29 cells. These results imply that at least two molecular pathways contribute to *C. difficile* mucosal adhesion. In addition to their direct roles encoding T4P and flagellar subunits, *pilA1* and *fliC* may contribute to regulating other factors relevant to mucosal adhesion.

## Introduction

*Clostridioides difficile*, a spore-forming, anaerobic, Gram-positive bacterium linked to colitis and antibiotic-associated diarrhea, has garnered significant scientific interest due to its public health impact (1). According to the CDC, *Clostridioides difficile* infects around 224,000 hospitalized patients and causes nearly 13,000 deaths annually in the United States (2). Infections primarily occur in hospitals following antibiotic treatments that disrupt the gut microbiome (3). This disruption enables toxin-producing *C. difficile* to proliferate, resulting in *C. difficile* infection (CDI) (4). As an obligate anaerobe, *C. difficile* vegetative cells cannot survive in aerobic environments outside a host (5). In response to environmental cues like nutrient limitation, quorum sensing, and other stress factors, *C. difficile* initiates sporulation to produce dormant spores capable of withstanding harsh conditions (5–7) on their own or as components of *C. difficile* biofilms (8).

For many organisms, adhesion to gastrointestinal cell surfaces is crucial for colonization and the production of virulence factors (9). Mucin proteins are heavily glycosylated and can be divided into those bound to the cell membrane and those which are secreted into extracellular space (although some membrane-bound mucins can be secreted through cleavage) (10, 11). Mucins, produced and secreted by goblet cells in the intestine, form a gel-like protective barrier that shields the intestinal epithelium from the gut microbiota (9, 12). In the colon, which is the primary site of *C. difficile* colonization (3), the mucus layer is often described as being divided into two layers, with an outer layer that serves as the site of bacterial colonization, and an inner, sterile layer (13, 14). Mucin proteins have distinct roles and those with known mechanisms from prior studies include MUC2, which supports the growth of commensal microbes, and MUC1, a membrane-bound mucin essential for defense against bacteria (13, 15, 16).

Flagella are crucial for bacterial swimming motility and promote gastrointestinal colonization in many bacterial species, acting as key virulence factors in early infection, as seen for GI pathogens *Escherichia coli*, *Salmonella Typhimurium*, *Helicobacter pylori* and *Vibrio anguillarum* (17–20). For *C. difficile,* flagella have been shown to play essential roles in adhesion to and colonization of the gut (21). Prior studies have shown that *C. difficile* mutant strains lacking the major structural protein, flagellin (FliC) and the flagellar cap (FliD) have reduced adhesion to mouse mucus (22) and exhibit a reduced capacity to adhere to the mouse cecum (22, 23). However naturally-occurring aflagellate clade V *C. difficile* strains have been shown to adhere to mucosal surfaces *ex vivo* (24).

The mechanisms by which *C. difficile* interacts with the mucus layer and underlying epithelium to promote colonization are not well understood, even though this association is known to occur. Our group and others have previously demonstrated that *C. difficile* is able to bind directly to mucins from human colonic cell lines and porcine gastric and colonic tissues when tested using *ex vivo* mucosal surfaces (24, 25). We demonstrated that *fliC* mutant strains had reduced adherence to gastrointestinal mucins compared to the parent strain. This finding suggested that interactions between host mucins and *C. difficile* flagella facilitate the initial host attachment of *C. difficile* to host cells and the production of mucus. However, as FliC is also known to regulate the expression of other genes in *C. difficile* (26), we could not exclude an indirect mechanism by which FliC could mediate mucin binding. We also showed that a mutant unable to produce the type IV pilus due to a mutation in *pilA1*, the gene encoding the major structural protein of the Type IV pilus, bound at higher levels than wild type bacteria; although, we could find no direct explanation for increased adherence from the loss of an adhesive appendage.

Previous work from Anjuwon-Foster *et al.* showed that motility and toxin production are regulated by phase variation in *C. difficile* strain R20291 (27). They discovered an invertible riboswitch upstream of the F3 gene cluster, which is essential for flagella formation (Fig. 1). To further characterize the effects of phase variation on *C. difficile* physiology and pathogenesis, the Tamayo group created small deletions in the invertible switch region to generate, *flg*-ON and *flg-*OFF mutants (28, 29). The *flg*-ON variant showed greater swimming motility and toxin production *in vitro*, and caused greater weight loss and a higher level of colonization in an *in vivo* mouse model. However, the *flg*-OFF variant showed greater lethality in a hamster model of infection. These findings suggest that phenotypic variants generated by flagellar phase-switching have distinct potentials for disease progression (29, 30).

**Figure 1.**
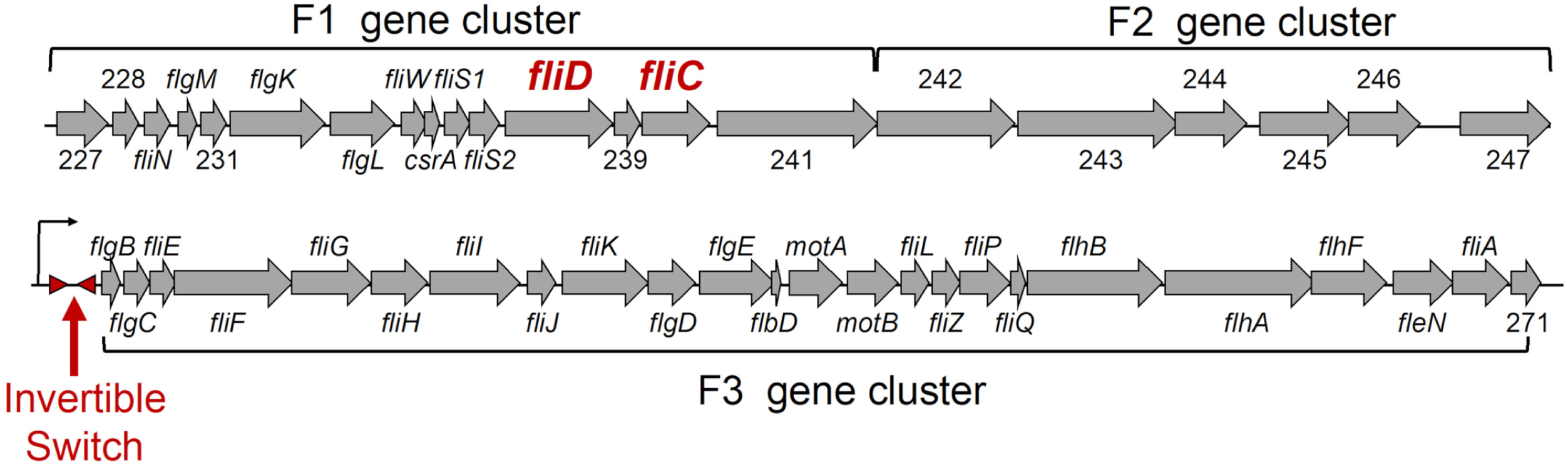
Flagellar operons in *C. difficile* strain R20291. Flagella production is governed by three major gene clusters in *C. difficile*, F1, F2, and F3. These gene clusters encode structural proteins, regulators, transporters, modification enzymes, and genes of unknown function (locus tags indicated by numbers). In *C. difficile* R20291, expression of the F3 gene cluster is regulated by an invertible regulatory switch that allows or inhibits gene expression. The presence of this invertible switch varies across strains of *C. difficile*.

To probe the potential roles played by flagella in mucosal attachment, this study examined how mutations to multiple flagellar genes (*fliC*, *fliD*), *pilA1* and phase variants *flg*-ON and *flg*-OFF in *C. difficile* R20291 impacted flagellation, macroscopic swimming motility, and adhesion to *ex vivo* mucosal surfaces. We found that, like the parent strain*, flg*-ON *and pilA1* had peritrichous flagella, while *fliC*, *fliD*, and *flg*-OFF were aflagellate; *pilA1* and *flg*-ON were hyperflagellated compared to the parent strain. We observed that flagella were required for macroscopic swarming motility, but hyper-flagellation did not increase motility. The *pilA1* mutant exhibited a significant increase in mucosal adhesion, whereas the *flg*-ON strain exhibited a minor decrease in binding compared to the wild type and all aflagellate strains showed significantly decreased binding. Similar roles for *fliC* in mucin adherence were seen in a cell culture-based model of mucin adherence. Measurements of transcription through qRT-PCR data indicate that hyper-flagellation in *pilA1* or *flg*-ON is not due to increased *fliC* expression and that *fliC* expression was significantly lower in both the hyperflagellated *flg*-ON and the aflagellate *flg*-OFF strain compared to the wild type (the *fliD* mutant had *fliC* transcription levels comparable to the parent strain). Taken together these results suggest that flagellation directly promotes adhesion, but that hyperflagellation alone is insufficient to promote strong binding, suggesting a role for other adhesins. There is a significant gap in understanding *C. difficile* colonization of the gut by *C. difficile*, particularly regarding its interactions with host cells and mucosa. Our findings here help to interpret previous reports of adhesion abnormalities in flagellin and pilin mutants, emphasizing the crucial role of flagellin in adherence to mucosal hydrogels.

## Materials and Methods

### Bacterial strains and growth conditions

Wild type and mutant strains of *Clostridioides difficile* R20291, a well-studied ribotype 027 clinical isolate, were used in this study (31). Wild type *C. difficile* R20291 as well as the *pilA1* and *fliC* mutant strains were created by Glen Armstrong (The University of Calgary) as previously described (24, 32). The R20291 *fliD* gene-interruption mutant was also created by Glen Armstrong using the CLOSTRON mutagenesis system as described for the CD630 *fliD* mutant in Dingle *et al.* (33). Phase-locked variants *flg*-ON and *flg*-OFF (16) were a generous gift from Rita Tamayo (University of North Carolina at Chapel Hill), as was her wild type R20291 strain. Brain Heart Infusion Supplemented (BHIS) Media was prepared from Difco BHI media supplemented with 5 g/L yeast extract and was used for all growth assays. BHIS media was prepared without cysteine supplementation to limit the potential for hydrogen sulfide production. For routine culture, BHIS agar plates were prepared with 1.5% w/v agar. All growth was performed at 37°C in an anaerobic chamber (Coy Laboratory Products) with an atmosphere of 90% nitrogen, 5% carbon dioxide, and 5% hydrogen. At the start of each experiment, *C. difficile* strains were inoculated from a glycerol stock onto pre-reduced BHIS agar plates. Details for subsequent growth steps unique to each assay are noted below.

### Preparation of artificial mucosal surfaces

Porcine colonic mucin extraction and purification were performed as previously described (24). Briefly, porcine colonic mucin, resuspended at a final concentration of 1 mg/mL in mucin coupling buffer (0.0308M K₂HPO₄, 0.0192M KH₂PO₄, 0.15M NaCl and 0.01 M EDTA) was used to coat glass coverslips (25 x 25 mm) with mucin using our previously published APTES-mediated coupling approach (24) with the following modifications. After modification with APTES and glutaraldehyde, each 25 x 25 mm glass coverslip was submerged in 3.5 mL of mucin solution in a 55 mm diameter polypropylene petri dish (Eisco labs) and incubated overnight on an orbital shaker at 4°C. To ensure even mucin adsorption, coverslips were turned over, returned to the mucin solution, and put back into the shaker the following day at 4°C for an additional four hours. Next, coverslips were washed with phosphate-buffered saline (PBS, 137 mM NaCl, 2.7 mM KCl, 10 mM Na_2_HPO_4_, 1.8 mM KH_2_PO_4_, pH 7.4) and exposed to UV light for sterilization. Mucin-coated coverslips were stored at 4°C in a coverslip holder (Sigma Aldrich) within a sterile container for future use (1).

### Bacterial adherence to mucosal surfaces

For all strains, five to ten colonies were inoculated into BHIS broth and grown anaerobically at 37°C until mid-exponential phase (Optical Density at 600 nm = 0.5, approximately 4-6 hours). Bacterial cells were centrifuged at 5000 x g for 5 minutes, decanted under anaerobic conditions, then resuspended in anaerobic phosphate-buffered saline (PBS) at an OD600=0.5. An aliquot of the cell suspension was removed for determination of colony forming units (CFU) through serial dilution and plating. 6 ml of cell suspensions were added to each well of sterile, six-well cell culture plates (Costar, Corning Incorporated) that contained mucin-coated glass coverslips that had been pre-reduced for 2 hours. Cells were incubated with coverslips for one hour at 37°C. Binding was also assessed to control glass coverslips that were treated as described above, save that the incubation was in mucin coupling buffer without added mucin. The coverslips were carefully rinsed four times with PBS to eliminate any non-adherent bacteria. Rinsing was performed by sequential dipping of coverslips into sterile, square polypropylene containers (4 cm^2^) that contained 40ml of PBS. To measure the adherent bacterial population, coverslips were then immersed in a 0.25% trypsin-EDTA solution (Gibco) at 37°C for 10 minutes. Trypsin activity was neutralized by adding two volumes of BHIS broth and levels of adherent cells were determined by serial dilution and plating of cells onto BHIS agar plates, which were incubated at 37°C for 24 hours to facilitate colony growth and enumeration. After 24 hours, the number of colonies per plate was counted and percent binding was calculated as the CFU/mL of adherent bacteria divided by the CFU/mL inoculum of bacteria.

### Bacterial Adherence to HT-29 and HT-29 MTX 2D cell culture

Authenticated HT-29 and HT-29 MTX-E12 cell lines were obtained from Sigma-Aldrich and cultured in 75-cm² plastic flasks (Fisher Scientific). HT-29 cells were maintained in Roswell Park Memorial Institute 1640 (RPMI 1640) medium with L-glutamine and sodium bicarbonate (Sigma-Aldrich), supplemented with 10% (v/v) fetal bovine serum (FBS), 1% penicillin-streptomycin solution (10,000 U/mL, Gibco), and 1% Amphotericin B (250 μg/mL, Gibco). HT-29 MTX-E12 cells were cultured in Dulbecco’s Modified Eagle Medium (DMEM; 4.5 g/L glucose and L-glutamine, without sodium pyruvate), supplemented with 10% (v/v) FBS, 1% penicillin-streptomycin solution (10,000 U/mL, Gibco), 1% Amphotericin B (250 μg/mL, Gibco), 1% MEM non-essential amino acids (100x, Gibco), and 1% GlutaMAX (100x, Gibco).

For use in the experiments, 12-well tissue culture plates (Fisher Scientific) were seeded with 100,000 cells/well. HT-29 and HT-29 MTX-E12 cells were used at passages 8–13 and 57–59, respectively. The culture medium was replaced three days after seeding and subsequently every two days for a total of 18 days. All cultures were maintained at 37°C in a humidified atmosphere of 5% CO₂ and 95% air. A 12-well plate containing confluent monolayers of HT-29 and HT-29 MTX-E12 cells was gently washed twice with pre-warmed, bicarbonate-free, phenol-red-free Hanks’ Balanced Salt Solution (HBSS). For both strains of *C. difficile* R20291 (wild-type and *fliC* mutant), five to ten colonies from BHIS agar plates (grown anaerobically at 37°C for 24 h) were inoculated into BHIS broth and cultured overnight under anaerobic conditions at 37°C. The overnight cultures were centrifuged at 4,700 × g for 8 min at 20°C, and the resulting bacterial pellets were resuspended in anaerobic HBSS in the original culture volume. A 1 mL aliquot of this bacterial suspension was anaerobically added to each well of the 12-well plate containing the washed monolayers. The plates were incubated anaerobically at 37°C for 2 h. Following incubation, supernatants were collected, serially diluted, and plated on BHIS agar to enumerate viable bacteria after the 2-hour incubation. To quantify bound bacteria, the remaining supernatant was discarded, and the wells were washed three times with HBSS to remove unbound bacteria. The monolayers were then scraped into 1 mL of HBSS, thoroughly mixed, and serially diluted before plating onto BHIS agar plates. After 24 h of anaerobic incubation at 37°C, colony-forming units (CFUs) were counted. The percentage of adherence was calculated as the CFU/mL of bound bacteria divided by the total CFU/mL of bound and viable bacteria after 2 hours of incubation, multiplied by 100. To determine bacterial attachment to plastic wells, the same procedure was conducted using 12-well plates without cell monolayers.

### Swimming motility assay

For swimming motility assays, BHIS agar plates were prepared with 0.3% w/v agar. *C. difficile* strains were grown in BHIS broth as described above, then three microliters of broth culture were spotted onto pre-reduced BHIS 0.3% agar plates. Following inoculation, plates were incubated anaerobically for 48 hours at 37°C in a partially sealed plastic container to prevent drying of the agar. The extent of bacterial swimming was assessed at 48 hours as described previously (34).

### Measurement of flagellation by TEM

The number of flagella on wild type and mutant *Clostridioides difficile* R20291 cells was quantified using transmission electron microscopy (TEM). As above, *C. difficile* strains were grown anaerobically in BHIS broth at 37°C for 4-6 hours until OD600 = 0.5. Bacterial cultures were fixed by combining with an equal volume of 2.5% glutaraldehyde in 100 mM cacodylate buffer, pH 7.4, and incubating at room temperature for one hour. Cells were washed twice in the same buffer. Samples were stored at 4°C until they were prepared for TEM examination at the UNL Center for Biotechnology’s Microscopy Research Core Facility (35). To set bacteria on TEM grids, a 30 ml droplet of bacterial suspension was placed on parafilm and transferred to a 200-mesh carbon-formvar coated copper grid by placing the grid, coated-side down, in contact with the samples for 30-60 seconds. After that, excess bacterial suspension was wicked away by touching a piece of filter paper to the edge of the grid surface and the samples were air-dried for 30 sec. The grid was placed with sample-side down onto a droplet of stain solution (1% uranyl acetate) for 30-60 seconds. The excess staining solution was wicked from the grid following the staining. The grids with sample were allowed to air-dry for 10 min before they were examined at 80kv under a Hitachi HT7800 TEM. Ten to fifteen TEM micrographs were obtained per strain at multiple magnifications with 7000x images used for quantification in Figure 2.

**Figure 2.**
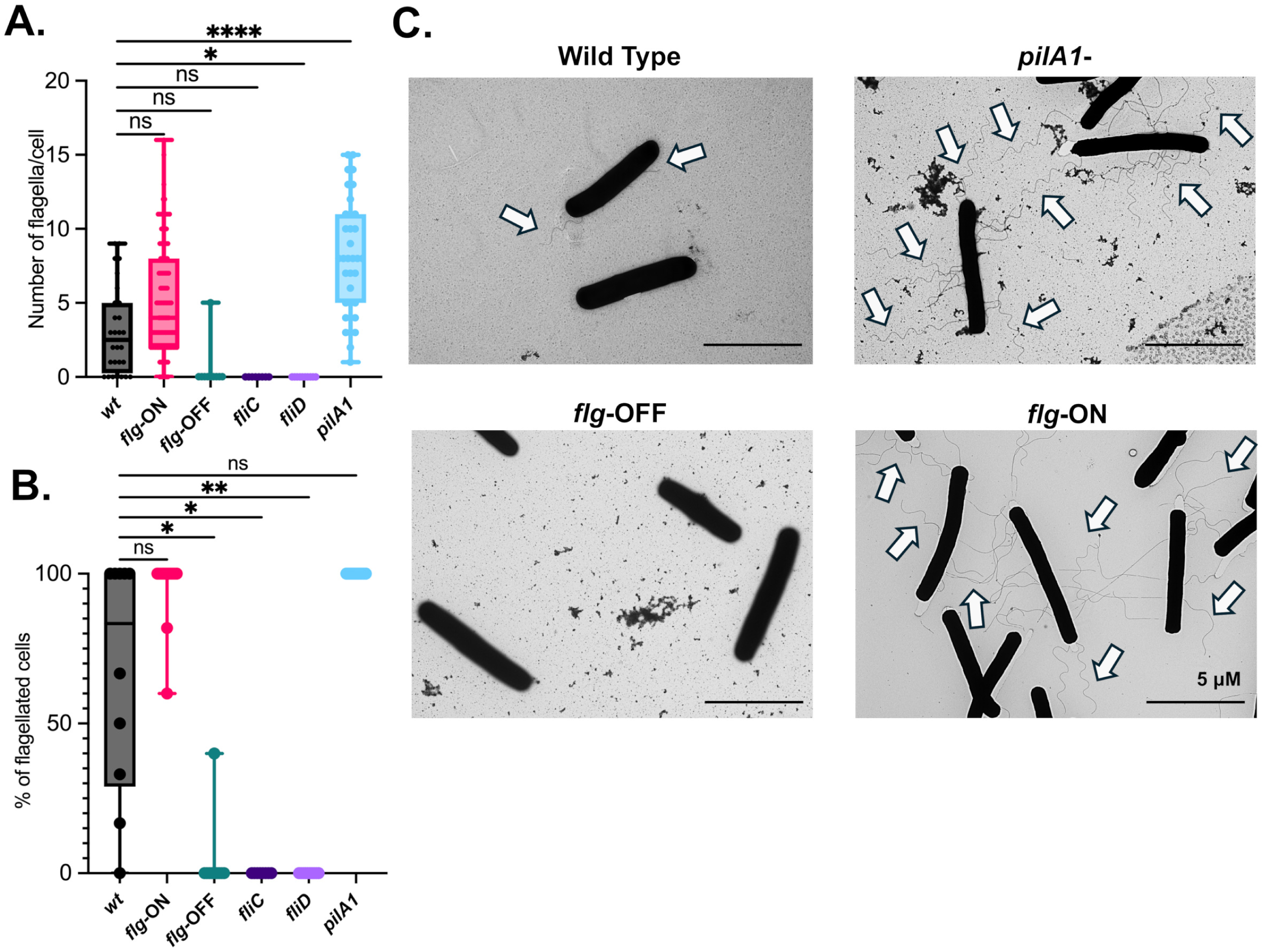
*flg-*ON and *pilA1* mutations increase the number of flagella per cell. (A) Representative transmission electron microscopy images of wild type, *pilA1, flg*-ON*, fliC, fliD and flg*-OFF strains. (B) The number of flagella per cell and (C) percent of flagellated cells were determined from enumeration of TEM images. Significance of differences from wild type were determined with Kruskal-Wallis testing with Dunn’s correction for multiple comparison. ****, p<0.0001; *, p<0.05, ns = not significant.

### fliC expression measurement through qRT-PCR

Bacteria were grown from frozen stocks on BHIS agar plates for 24 hours. 10 colonies were added to 14 ml of BHIS broth and incubated at 37°C until a cell growth of OD_600_∼0.5 was reached. Cells were then mixed with an equal volume of ice-cold methanol to fix cells and centrifuged for 10 minutes at 3000 × g at 4°C to pellet cells. Pellets were washed in 500 µl of Tris EDTA (10 mM Tris-HCl, 1 mM EDTA, pH 7.5), centrifuged for three minutes at 4000 × g at 4°C, and resuspended in 1 mL Buffer RLT obtained from the Qiagen RNeasy kit. 0.1 mm silica-glass beads were added to samples and cells were disrupted by bead beating for one minute. The supernatant was carefully removed, and 1 volume of 70% ethanol was added to the lysate and mixed well by pipetting. RNA extraction then proceeded according to Qiagen protocol with optional DNase I digestion (Qiagen). RNA concentration was determined by UV spectrometry and quality was assessed by stable RNA (16S, 23S rRNA) band integrity following gel electrophoresis. qRT-PCR was performed in two steps. In the initial reverse transcription step, 5 mg of total RNA was incubated with 1 µL of 50 mM random hexamers (Invitrogen), and 1 µL of 10 mM dNTP mix (Invitrogen) in a total volume of 12 µL at 65°C for 5 min. Reverse transcription was then performed with Superscript II Reverse Transcriptase (Invitrogen). specifically, 4 µL of 5X First-strand Buffer, 2 µL of 0.1 mM DTT, and 1 µL of RNase OUT Recombinant Ribonuclease Inhibitor (Invitrogen) were added to each reaction, which was then incubated at 25°C for 2 min. 1 µL of Superscript II reverse transcriptase was then added to each reaction and reactions were incubated at 25°C for 10 min, 42°C or 50 min, and 70° for 15 min. 1 µL of Ribonuclease H (Invitrogen) was then added to each tube to degrade RNA/DNA hybrids before initiating qPCR. qPCR was performed using primers designed to amplify a section of the *fliC* gene (*fliC* forward: GCA CAA AGT AAG TCT ATG G; *fliC* reverse: CAG ATA TAC CAT CTT GAA CG) as well as the housekeeping gene (*rpoA*) that encodes RNA polymerase that encodes RNA polymerase (*rpoA* forward:CAT GCT CTA TCA CAG GTG CAG;*rpoA* reverse: CA ACT CTT GTG TTT TCC ACA). Duplicate qPCR reactions were set up for each reverse transcribed product.

20 µL reactions contained 1 µL of reverse-transcribed DNA, 300 nM forward and reverse primers, and 1X Power SYBR Green PCR mix (Thermo Fisher Scientific). Amplification was performed in a QuantStudio 3 PCR machine (Thermo Fisher Scientific) with an initial denaturation step for 10 min at 95°C, followed by 40 cycles of a 15-sec denaturation step at 95°C followed by a 60-sec annealing and elongation step at 60°C. Relative levels of *fliC* were determined through the ΔΔC_t_ method with the average of wild-type replicates used as a baseline for expression.

### Analysis of data and evaluation of statistical significance

Data were visualized and the statistical significance was determined with GraphPad Prism version (10.0.2) (Cambridge, MA). Tests performed in each experiment are indicated in the figure legends. Significance values less than p<0.05 are reported.

## Results

### Disruption of pilA1 and loss of phase variation in F3 operon impact flagellation

The flagellum of *C. difficile* is a highly studied component that contributes to colonization (33). In a previous study, we demonstrated that the major structural protein of flagella, FliC, contributed to mucin adhesion. While *fliC* mutant cells exhibited reduced adherence to mucin hydrogels compared to wild type cells, *pilA1* mutants, which lack the major structural protein of type IV pili, exhibited increased binding to mucosal surfaces. We hypothesized that this increased mucosal adhesion may be due to increased flagellation in a *pilA1* mutant. To test this hypothesis, TEM images were collected from wild type and mutant *R*20291 strains and the number of flagella per cell (Fig. 2a) and the proportion of flagellated cells (Fig. 2b) were measured. Previous studies of *C. difficile* R20291 identified two primary lineages which display significant phenotypic variations (31). Germane to this study, CRG0825 strains show peritrichous flagellation while CRG2021 strains show a single polar flagellum. Our wild-type strain was peritrichously flagellated, with a mean number of flagella per cell of 3.1 ± 3.0. However, not all cells (67 ± 39%) were flagellated. As we expected, the *fliC* mutant strain lacked flagella, as did cells with mutations in *fliD,* which encodes the flagellar cap. The percentage of flagellated cells and the mean number of flagella per cell both rise in the *pilA1* mutant to 100% and 8.1 ± 3.9 respectively, consistent with our hypothesis that the *pilA1* mutant is more highly flagellated.

We also assessed the number of flagella per cell and percent flagellation in cells where phase variation of the F3 operon was locked in the “ON” or “OFF” position due to a three-nucleotide deletion in the invertible switch (36). Consistent with prior observations (27)*, flg*-OFF mutants were primarily aflagellate. In contrast, *flg*-ON mutants were peritrichously flagellated, with the mean number of flagella per cell 5.2 ± 3.7 (Fig. 2a) and percentage of flagellated cells 96 ± 11% (Fig 2b), significantly higher than wild type cells. Based on these results, we conclude that both *pilA1* and *flg*-ON mutants are hyperflagellated, whereas *fliC, fliD, and flg*-OFF mutants are primarily aflagellate.

### Macroscopic swimming motility is unaffected by hyper-flagellation

To investigate the relationship between hyper- and hypo-flagellation and swimming motility, we measured macroscopic swimming motility for the strains described above. Swimming motility was assessed by measuring the migration of cells through 0.3% BHIS agar over a 48-hour incubation. We observed that the mean distance moved for wild type cells was 16 ± 0.71 mm. *flg*-ON and *pilA1* mutants exhibited comparable motility zones of 17±1.7 mm and 17±1.8 mm respectively. Although hyper-flagellation did not enhance macroscopic swimming motility, the *flg*-OFF, *fliC* and *fliD* mutants exhibited significantly less motility than wild type with motility zones of 5.6±0.25 mm, 5.3±0.29 mm, and 5.1±0.25 mm, respectively. Overall, the results support a binary model of swimming motility over this timescale with the wild type, *pilA1*, and *flg*-ON strains showing similar motility, while the *fliC, fliD*, and *flg*-OFF mutants were non-motile.

### Flagellar mutants are defective in adhesion to mucosal hydrogels

In our previous study, we observed that the *fliC* mutant exhibited impaired binding to mucosal hydrogels formed on glass surfaces, while the *pilA1* mutant showed a modest increase in binding (24). Given that *fliC* is a pleiotropic regulator that potentially regulates surface adhesins, we evaluated mucosal adhesion in other mutant strains (f*liD, flg*-ON*, and flg*-OFF) with altered levels of flagellation to determine whether variation in other flagellar components might impact mucosal binding (37). For wild type R20291 cells, we observed that 0.14 ± 0.099% of cells bound to mucin after a 1-hour incubation (Figure 4). In line with our previous results, *pilA1* showed increased binding at 1.5± 2.0 %, whereas the *fliC* mutant showed essentially no binding (0.0060 ± 0.0046%). Similarly, *fliD* (0.018 ± 0.017%), and *flg-*OFF (0.023 ± 0.037%) demonstrated reduced binding compared to the wild type. However, in contrast to the *pilA1* mutant, *flg-*ON (0.052 ±0.037 %) showed a slight reduction in binding relative to the parent strain rather than increased binding as hypothesized.

**Figure 3.**
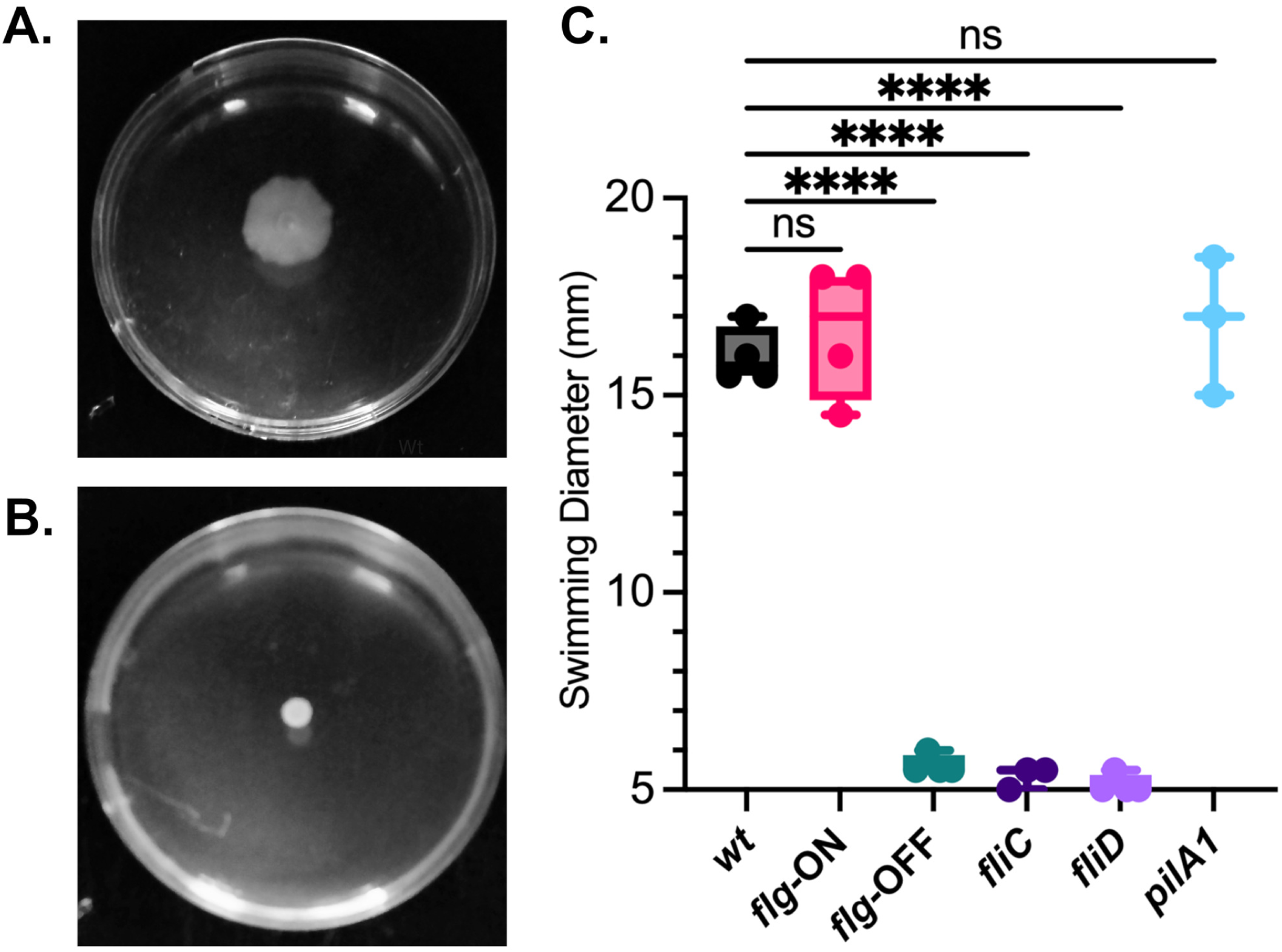
Differences in macroscopic swimming motility between R20291 strains. Flagellar motility was measured on 0.3% BHIS agar plates. (A). Representative images of wild type, *pilA1, Flg-On, fliC, fliD and Flg-Off*. (B) Motility was enumerated and plotted with significance of differences from wild type determined by Brown Forsythe and Welch ANOVA with Dunnet T3 correction for multiple comparison. ****, p<0.0001; ns = not significant.

**Figure 4.**
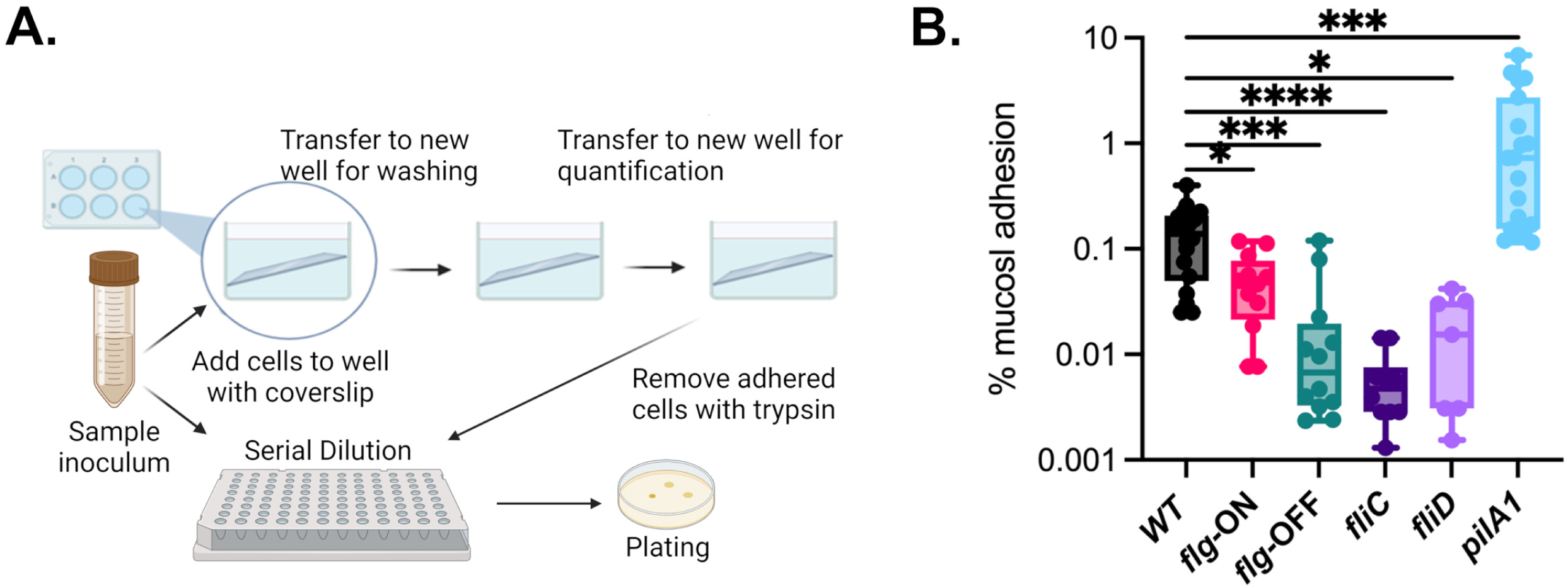
*C. difficile* binds specifically to mucin hydrogels *in vitro*. (A) Overview of approach to quantify mucin-specific binding through selective plating. (B) Comparison of wild type and mutant *C. difficile* R20291 adhered to mucin-hydrogels. Significance of differences from wild type was determined by Brown-Forsythe and Welch ANOVA test with Dunnet T3 correction for multiple comparisons. ****, p<0.0001; ***, p<.001; **, p<0.01; *, p<0.05; ns = not significant.

### Loss of flagellation decreases adherence to mucin-secreting cells in 2D culture

Previous studies measuring adherence by *C. difficile* to 2D Caco-2 cell monolayers showed strain-dependent defects in binding for *fliC* mutants; a defect was found in the R20291 background (24), but not in CD630 (38). Because cells in 2D culture can both secret mucins and/or express membrane-bound mucins on their surfaces, this defect could also be explained as a defect in mucosal adhesion. To test hypothesis, we compared the adhesion of *C. difficile* R20291 wild type and *fliC* bacteria to monolayers of HT-29 and HT29-MTX cells. The HT-29 cells line was derived from a colorectal adenocarcinoma (39) and HT-29 MTX cells were derived as a stable subpopulation from HT-29 cells after treatment with methotrexate (40). This model has been used previously to measure the impact of mucus-secretion on adherence by anaerobic bacteria (41). We also previously found that the R20291 *fliC* mutant has a defect in adhesion to mucosal hydrogels formed with mucins purified from HT-29 MTX cells compared to the parent strain (24). Consistent with previous observations, we found that adherence of R20291 wt cells to HT29-MTX cells (67 ± 5.5%) was >180-fold higher than adherence to HT29 cells (0.37 ± 0.038%) (FIgure 5). Binding of the *fliC* mutant to HT-29 MTX cells (32 ± 2.0%) was significantly lower than wt cells, whereas there were no significant differences in binding to HT29 cells (0.53 ±0.052%) between wt and *fliC* mutants. These results provide further support for the importance of the mucin layer as a site for *C. difficile* attachment and demonstrate that *fliC* contributes to this adhesion.

### Hyper-flagellation is not mediated by increased expression of fliC

Because of the potential regulatory role of *fliC*, the decrease in adhesion of the *fliC* mutant (and the increased mucosal adhesion of the *pilA1* mutant) could be driven either by direct adhesion of flagella or regulation of other adhesins by FliC. We used qRT-PCR to compare *fliC* transcription levels across *C. difficile* strains. We grew R20291 wild type and the *pilA1*, *flg*-ON, *flg*-OFF and *fliD* mutant strains and measured *fliC* expression by qRT-PCR, normalizing the results to the average expression level in the wild type strain. Transcription of *fliC* varied across replicate samples collected from wild type strains, with a mean level of log_2_ normalized expression of 0 ± 1.5 (Figure 6). No significant differences in *fliC* expression were observed between *fliD* (1.3 ± 1.4) and *pilA1* (0.9 ± 0.8) and the parent strain. However, both *flg*-ON (−3.7 ± 1.1) and *flg*-OFF (−2.3 ± 0.38) exhibited significantly lower *fliC* expression compared to wild type. Because *flg*-ON and *flg*-OFF were obtained from the Tamayo lab and may be of a different genetic lineage, we also compared *fliC* expression in our R20291 wild type strain to the R20291 wild type strain obtained directly from Rita Tamayo’s lab, wild type_RT_. However, no significant differences in *fliC* expression were observed between the two wild type strains.

**Figure 5.**
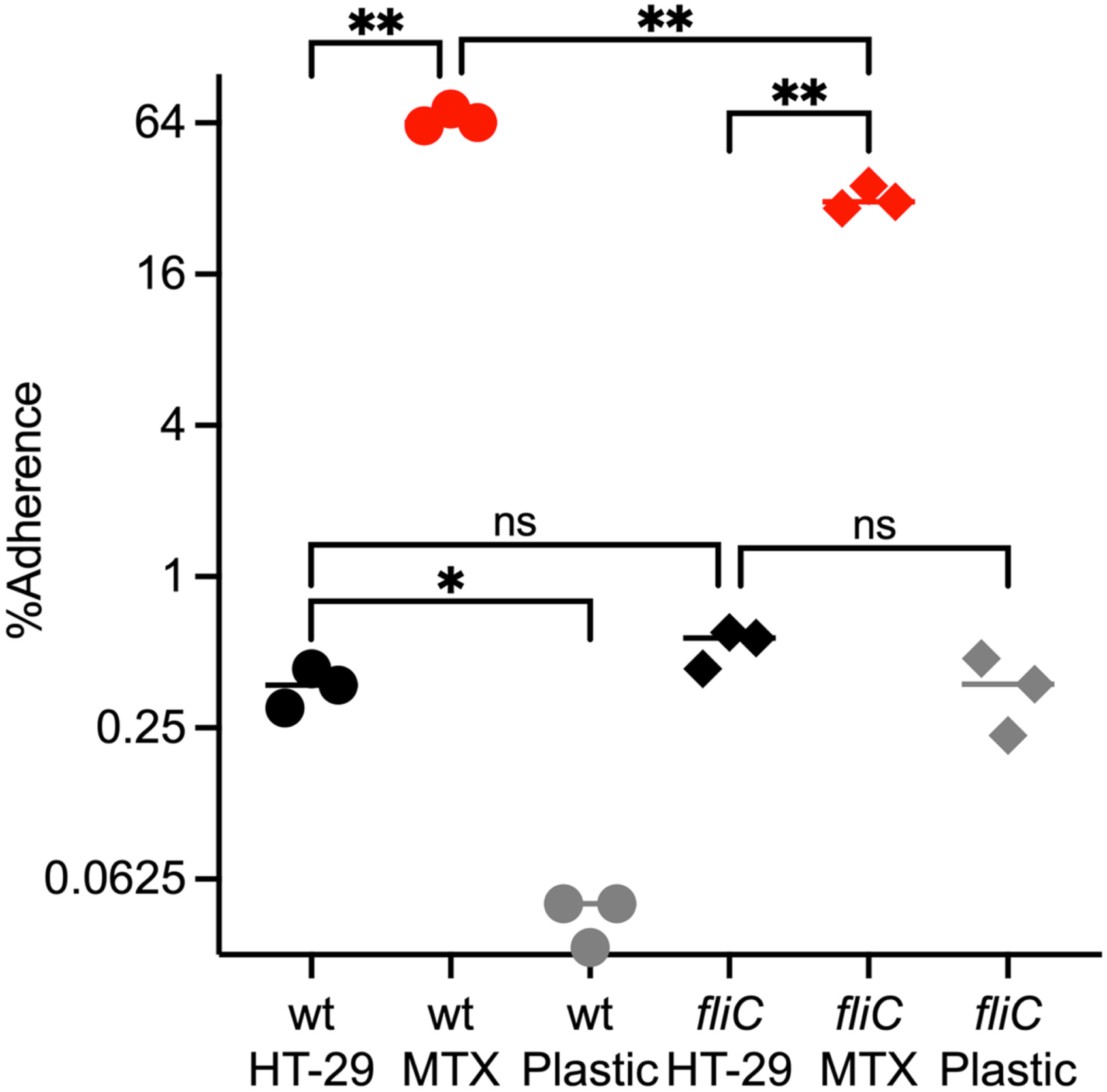
Adherence to HT-29 and HT-29 MTX cells. Percentage adherence) is shown for R20291 wild type (circles) and *fliC* (diamonds) to HT-29 cells (black), HT-29 MTX cells (red) and the plastic surfaces of empty wells (grey). Statistical significance was determined by Brown-Forsythe and Welch ANOVA with Dunnett T3 correction for multiple comparisons. *, p<0.05, **, p<0.01; ns = not significant.

**Figure 6.**
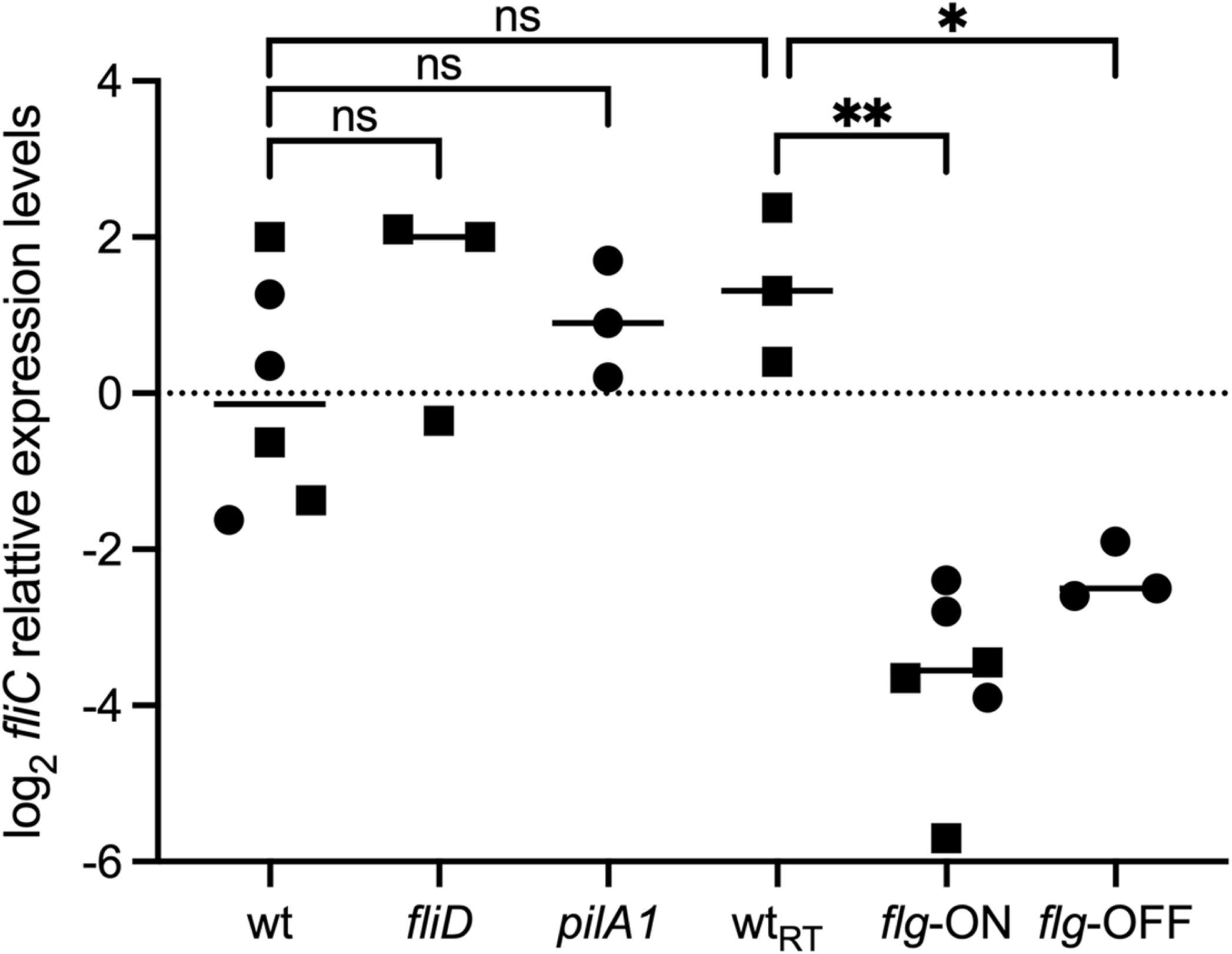
Impacts of mutations on *fliC* gene expression. Comparisons of relative levels of *fliC* gene expression between R20291 wild type (received from from either Glen Armstrong’s group (wt) or Rita Tamayo’s group (wt_RT_), *pilA1, fliC, fliD* (all derived from wt)*, flg*-ON*, and flg*-OFF (derived from wt_RT_) through qRT-PCR. Squares and circles indicate different experiments. Statistical significance was determined by Brown-Forsythe and Welch ANOVA with Dunnett T3 correction for multiple comparisons.

## Discussion

The high rates of recurrence, the emergence of hypervirulent strains, and the ongoing increase in the incidence of *C. difficile* underscore the necessity of identifying pharmacological targets and developing innovative therapeutic approaches to treat *C. difficile* (42). However the identification of specific molecular mechanisms critical for *C. difficile* colonization is complicated by the interplay between host-pathogen and inter-microbial interactions. (42). The gut microbiota disruption created by antibiotics creates conditions favorable for *C. difficile* colonization and toxin production (43–50). Although it is known that *C. difficile* exploits an intestinal environment after disruption of the commensal microbiota, the precise mechanism of these host-microbe and inter-microbial interactions is unknown (51). Multiple studies have shown that *C. difficile* adheres to host-derived cells in 2D culture (52–59); more recently, strong interactions have been observed *C. difficile* adherence to mucus both *in vivo* (25) and *ex vivo* (24). Competition between microbes for host mucins as attachment sites and microbial degradation of host mucins as nutrient sources are potential factors influencing the competition between *C. difficile* and the commensal microbiome. These interactions with host mucosa could provide another avenue for commensal microbes to contribute to *C. difficile* colonization.

Previous studies of *C. difficile* R20291 have shown that flagellin is a key component of *C. difficile* adherence and persistence (23, 60), although similar experiments in CD630 have not seen equivalent effects (38). In addition to serving as the major structural component of flagella, FliC is also a pleiotropic regulator, with activities related to colonization, virulence, and toxin gene expression (22, 37, 61–63). Expression of *fliC,* toxin synthesis, bacterial movement, colonization, and pathogenicity are intricately regulated through a network that includes the global regulator CsrA (carbon storage regulator protein A). Previous studies have shown that FliW influences *fliC* expression by obstructing CsrA-mediated post-transcriptional regulation (64). Additionally, FliW has been shown to suppress CsrA *in vivo*, which may have detrimental effects on *C. difficile* pathogenicity (64). Additional factors that regulate FliW, CsrA, and FliC, as well as the function of FliW in *C. difficile* are still unknown, even though the critical functions of CsrA in flagellin synthesis and flagellin homeostasis have been investigated in other bacteria (65). Further, flagellum and toxin expressions are linked through the flagellar alternative sigma factor, SigD, which is encoded in the F3 operon (66).

Phase variation of the F3 operon also contributes to regulation of flagella and toxin expression (36, 67). However, the role that this regulation plays in pathogenesis is complex. While phase-locked *flg-*ON mutants accumulated more toxins than the phase-locked *flg-*OFF mutants in a hamster model of acute *C. difficile* infection, they were not substantially different in their capacity to induce acute illness symptoms. In mice, *flg*-ON mutants showed both greater persistence and greater morbidity (36), but the *flg*-OFF mutant colonized at levels intermediate between wild type and *flg*-ON. In *in vitro* assays, *flg*-ON mutants have previously been shown to be flagellated and motile, unlike their *flg*-OFF counterparts (27).

Recently we demonstrated that both O-linked glycosylation of host mucins and *C. difficile fliC* were required for mucin adherence by *C. difficile* (*24*). While we hypothesized that direct interactions between flagella and mucins were responsible for this adherence mechanism, an alternative hypothesis is that the synthesis of downstream lectin-like adhesins is controlled by the expression of *C. difficile* FliC. Here, we have demonstrated that mutants lacking flagella but not FliC (*fliD*, *flg*-OFF) also exhibited reduced mucin binding. Conversely, while both *pilA1* and *flg-*ON mutants were hyper-flagellated, we observed that only *pilA1* mutants exhibited increased mucin binding. These results indicate that while flagellation is one factor contributing to mucin adherence, other factors likely contribute to mucin adherence in the *pilA1* mutant. Similarly, despite being hyper-flagellated, *pilA1* mutants show similar *fliC* expression levels to the parent strain, indicating that hyper-flagellation is not regulated through transcription of *fliC.* Further, the *flg*-ON and *flg*-OFF mutant strains show lower levels of *fliC* expression compared with the parent strain. This result differs from what was previously observed for naturally-occurring R20291 phase variants (27), but is similar to a comparison of phase-variants in CD630 which showed both flagellate and aflagellate derivatives showed lower *fliC* expression than the parent strain (67). This result suggests that *fliC* expression may be regulated by expression of the F3 operon, with both under and over-expression of other flagellar components leading to feedback regulation through an unknown mechanism.

Our observation that *fliC* contributes to mucin adherence was not limited to binding to synthetic mucosal hydrogels, as we also observed defects in *fliC* adherence to HT29-MTX cells, a 2D cell culture model that was previously shown to support mucin-specific adherence of *Clostridium perfringens* (*41*). In contrast, there were no significant differences in binding to HT29 cells between wt and *fliC* cells. A previous report has also shown an approximately 5-fold reduction in binding of a *fliC* strain to Caco-2 monolayers compared to wild-type (62). Caco-2 cells produce MUC1, MUC3, MUC4 and MUC5A/C (68), whereas HT29-MTX cells produce high levels of MU5A/C and low levels MUC2 (69). Differences in the effects of *fliC* mutants in these assays could possibly be due to the types of mucins produced or variations between *C. difficile* strains. Further studies are needed to explore these differences more fully. We also observed significantly higher non-specific binding of *fliC* cells to plastic compared to wild type cells. The significance of higher non-specific binding by *fliC* mutants is unclear, although these results do emphasize the importance of negative controls in these types of *C. difficile* studies due to high levels of plastic adherence.

Based on these combined results, we conclude that flagellation is not generally limited or controlled by levels of *fliC* expression but by other factors (likely including *fliD* expression and or other flagellin genes) and that flagellar adhesion alone cannot explain mucosal adherence by *C. difficile*. Instead, these findings suggest that *C. difficile* mucosal adherence is influenced by at least two molecular mechanisms, flagella-mediated binding and another adherence factor that is upregulated in *pilA1* mutants (8). Further studies are needed to identify this potential second adherence factor.

